# Cophylogeny and specificity between cryptic coral species (*Pocillopora* spp.) at Mo’orea and their symbionts (Symbiodiniaceae)

**DOI:** 10.1101/2022.03.02.482706

**Authors:** Erika C. Johnston, Ross Cunning, Scott C. Burgess

**Affiliations:** Department of Biological Science, Florida State University, 319 Stadium Drive, Tallahassee, FL, 32306-4296, USA; Daniel P. Haerther Center for Conservation and Research, John G. Shedd Aquarium, Chicago, IL, USA

**Keywords:** Coevolution, cospeciation, holobiont, mutualism, niche differences

## Abstract

The congruence between phylogenies of tightly associated groups of organisms (cophylogeny) reflects evolutionary links between ecologically important interactions. However, despite being a classic example of an obligate symbiosis, tests of cophylogeny between scleractinian corals and their photosynthetic algal symbionts have been hampered in the past because both corals and algae contain genetically unresolved and morphologically cryptic species. Here, we studied co-occurring, cryptic *Pocillopora* species from Mo’orea, French Polynesia, that differ in their relative abundance across depth. We constructed new phylogenies of the host *Pocillopora* (using genomic loci, complete mitochondrial genomes, and thousands of single nucleotide polymorphisms) and their Symbiodiniaceae symbionts (using ITS2 and psbA^ncr^ markers) and tested for cophylogeny. The analysis supported the presence of five *Pocillopora* species on the fore-reef at Mo’orea that mostly hosted either *Cladocopium latusorum* or *C. pacificum.* Only *Pocillopora* species hosting *C. latusorum,* and that have similar relative abundances across depths, also hosted taxa from *Symbiodinium* and *Durusdinium.* In general, the *Cladocopium* phylogeny mirrored the *Pocillopora* phylogeny. Within *Cladocopium* species, lineages also differed in their associations with *Pocillopora* haplotypes, except those showing evidence of nuclear introgression, and with depth in the two most common *Pocillopora* species. We also found evidence for a new *Pocillopora* species (haplotype 10), that has so far only been sampled from French Polynesia, that warrants formal identification. The linked phylogenies of these *Pocillopora* and *Cladocopium* species and lineages suggest that symbiont speciation is driven by niche diversification in the host, but there is still evidence for symbiont flexibility in rare cases.

## Introduction

For organisms that form symbiotic partnerships with microbes, the identity and composition of symbionts is often critical to the performance and function of the holobiont (the host and its symbionts) (Rohwer et al. 2002; Roughgarden et al. 2018; Van Oppen and Medina 2020). Especially when host-symbiont partnerships are obligate, variation among host individuals and species in their symbiont identity may explain some of the variation in their response to environmental gradients or disturbances (Innis et al. 2018; del Campo et al. 2020; Abbott et al. 2021). However, the extent to which the interaction between host and symbiont itself evolves remains unclear in many cases. Therefore, identifying host-symbiont specificity, and the extent to which host and symbiont lineages co-diversify through shared evolutionary histories, is necessary for understanding the ecological and evolutionary dynamics of the holobiont, and the degree to which symbioses promote or restrict adaptation to climate change (Takiya et al. 2006; Compant et al. 2010; Kaltenpoth et al. 2014; Seah et al. 2017)

Most scleractinian corals form obligate symbiotic relationships with photosynthetic dinoflagellate algae in the family Symbiodiniaceae. In some corals, symbiont flexibility and the environment often play a substantial role in the composition of Symbiodiniaceae hosted (Putnam et al. 2012; Cunning et al. 2015; Boulotte et al. 2016; Quigley et al. 2019). However, there is also evidence for some level of host-symbiont specificity (LaJeunesse and Thornhill 2011; Parkinson and Baums 2014; Forsman et al. 2020), though direct evidence for concordant evolutionary histories is often considered lacking (Rowan and Powers 1991; Van Oppen and Medina 2020).

The degree of host – symbiont specificity, and the potential for coevolution, should relate to the mode of symbiont transmission (Goodnight 2000; Baird et al. 2009, 2021; Zeng et al. 2017). Most (~85%) scleractinian coral species obtain their symbionts horizontally (i.e., acquired from the environment), which is common in species that spawn gametes with external fertilization (broadcast spawning) (Baird et al. 2009). These coral species are expected to host greater symbiont diversity, and necessarily host algal species that are able to live outside the coral (Fujise et al. 2021), reducing specificity and decoupling their evolutionary histories. In contrast, other coral species obtain their symbionts vertically (i.e., from parent to offspring; (Hirose et al. 2000; Hirose and Hidaka 2006). Vertical transmission is more common in coral species with internal fertilization, where the embryo develops within the polyp before release as a motile planula larva (brooders) (Baird et al. 2009). Coral species with vertical transmission are expected to host lower algal diversity (Bongaerts et al. 2015), and algal species that are highly host specialized and often unable to live and propagate outside the coral host (Fujise et al. 2021). As a result, partnerships with hosts that vertically transmit algal symbionts are expected to be more stable over time, facilitating coadaptation of partners as a result of the shared reproductive fate of both partners (Fisher et al. 2017). Therefore, especially in species with vertical transmission, the main cause of symbiont speciation is thought to be niche diversification provided by speciation of the host, since the host provides the habitat that modulates natural selection on the symbiont (Thornhill et al. 2014). Nevertheless, there are few direct tests of this hypothesis (Lewis et al. 2019; Turnham et al. 2021).

Estimates of cophyologeny, or the concordance between the phylogenies of hosts and symbionts, provide a means to identify host – symbiont specificity and shared evolutionary histories. Congruence between the phylogenies of coral hosts and algal symbionts may be the product of coevolution or co-speciation. Strictly speaking, coevolution involves reciprocity; that is, evolutionary change in one species in response to the traits of a second species, followed by evolutionary response by the second species to the change in the first species (Janzen 1980; Thompson 1994). Co-speciation is concordant patterns of speciation that do not necessarily involve reciprocity, such as speciation of the coral host and subsequent tracking and diversification of the algal symbionts (Lewis et al. 2019), or shared biogeography and similar responses to common environments (Nuismer et al. 2010; Althoff et al. 2012). However, cophylogeny between corals and their photosynthetic algal symbionts has not yet been definitively tested (LaJeunesse et al. 2010; Pinzon and Lajeunesse 2011), though there is recent evidence for short-term (~3 MYA) co-diversification between corals and algal symbionts (Turnham et al. 2021), and cophylogeny between corals and bacterial symbionts (Pollock et al 2018).

Estimates of cophyologeny are important because several factors could limit the concordance between host and symbiont phylogeny when it is expected to be high (i.e., when there is vertical transmission). Some coral species with vertical transmission appear to exhibit a mixed mode of symbiont acquisition, allowing larvae to associate with symbionts that are not detected in the maternal colony (Quigley et al. 2018). A mixed mode of symbiont transmission may allow juveniles to have greater flexibility in different environments than strictly vertical transmission. Even if symbionts are transmitted vertically, an individual can host a diverse community of ‘background’ symbionts, even when there is often only one taxa detected, that are transitory and may depend on environmental conditions (Lee et al. 2016; Rouzé et al. 2019; Kriefall et al. 2022; Strader et al. 2022). As a result, even in species with vertical transmission, the relationships between scleractinian host and symbiont can be dynamic over ecological timescales (Reich et al. 2017; Quigley et al. 2019). Certain species of algal symbionts may share evolutionary histories with the coral host, while other species within the same host clade may not. Identifying associations between host and symbiont that are maintained in different habitats and in response to environmental stressors is therefore necessary for confidently detecting coevolutionary relationships, as well as generating new hypotheses for what drives these associations.

Tests of cophylogeny between corals and their algal symbionts have been hampered in the past by unresolved taxonomy and phylogenetic relationships within both the coral host and the algal symbiont. Both the coral host and the symbiont contain morphologically-cryptic species, making it difficult to identify associations without rigorous genomic information. In particular, the coral genus *Pocillopora* is notorious for containing multiple cryptic species because of morphological plasticity and overlapping morphological phenotypes (Pinzón et al. 2013; Marti-Puig et al. 2014; Paz-García et al. 2015). Despite being a well-studied coral genus (Flot and Tillier 2004; Flot et al. 2008; Schmidt-Roach et al. 2014; Gélin et al. 2017; Johnston et al. 2017; Oury et al. 2021), with a monophyletic radiation approximately 3MY old (Johnston et al. 2017), the identification and placement of some mitochondrial lineages is not yet fully resolved. Similarly, Symbiodiniaceae, the dinoflagellates that form the symbiotic partnership with scleractinians, have recently received taxonomic revision at the family and genus level (LaJeunesse et al. 2018), and the extent to which variation in genetic sequences reflects species is only just beginning to be clarified (Thornhill et al. 2014; Lewis et al. 2019; Turnham et al. 2021).

Here, we investigate *Pocilloporα*–Symbiodiniaceae associations in multiple co-occurring cryptic *Pocillopora* species that differ in their responses to thermal stress and in their niche space across depths at Mo’orea, French Polynesia (Burgess et al. 2021; Johnston et al. 2021). Pocilloporid corals are a particularly interesting group because colonies that broadcast spawn actually transmit their symbionts vertically in their eggs (Hirose et al. 2000; Schmidt-Roach et al. 2012; Massé et al. 2013; Johnston et al. 2020), unlike many other broadcast spawning corals that do not transmit their symbionts to offspring (Baird et al. 2009). We used multiple genomic data sets to reconstruct phylogenetic relationships and determine the placement of mitochondrial haplotypes that were not included in previous *Pocillopora* phylogenomic analyses (Johnston et al. 2017). We also assessed Symbiodiniaceae relationships using sequence variation in both ITS2 and psbA^ncr^ markers in order to determine whether there is support for their cophylogeny, or multiple colonization events of distantly related Symbiodiniaceae into *Pocillopora.* Given the recent evidence of co-diversification described for some *Pocillopora* lineages with *Cladocopium* ((Turnham et al. 2021), we hypothesize that there will be specific *Pocillopora* - *Cladocopium* associations that are not likely to differ across depths, reflecting co-speciation.

## Methods

### Pocillopora *phylogenetic analyses*

#### Taxon sampling

Tissue samples from 44 *Pocillopora* colonies were collected using SCUBA in August 2019 from six sites (LTER 1 – 6) and three depths (5, 10, and 20 m) around Mo’orea, French Polynesia (Supplemental table 1). The samples contained nine mitochondrial open reading frame (mtORF) haplotypes including haplotype 1a, 2, 3a, 3b, 3f, 3h, 8a, 10, 11. Haplotype 1a contains two species: *P. meandrina* and *P. grandis* (Johnston et al. 2017). While colonies of the same clone are well known in *P. acuta* (haplotype 5) (e.g., Strader et al. (2022)), we did not sample clones. mtORF identification was carried out as described in Johnston et al. (2021) with haplotype numbering following Forsman et al. (2013) and Pinzón et al. (2013). The species names associated with certain haplotypes are only nominal names based on the morphospecies type (based on Veron and Stafford-Smith 2000) originally assigned to the first description of the haplotype in Johnston et al. (2021), Forsman et al. (2013) and Pinzón et al. (2013). Tissue samples were stored in salt-saturated DMSO (dimethyl sulfoxide) buffer (Gaither et al. 2011) until DNA was extracted. Specimens are stored at Florida State University.

#### DNA extraction and quantification

Genomic DNA was extracted from tissue using the OMEGA (BIO-TEK) E.Z.N.A. Tissue DNA Kit. Three elutions (50, 100, 100 μl) were collected in HPLC grade H_2_O. Extractions were inspected on a 1% agarose gel. All three elutions were combined and DNA was speedvac concentrated. Extractions were quantified using the Qubit dsDNA HS Assay Kit with the Qubit Fluorometer (ThermoFisher SCIENTIFIC).

#### Library preparation and sequencing

ezRAD libraries (Toonen et al. 2013) were generated following Knapp et al. (2016) as in Johnston et al. (2017). Briefly, samples were digested with the isoschizomer restriction enzymes *MboI* and *Sau3AI* (New England BioLab), which cleave at GATC cut sites, and libraries were generated with the KAPA HyperPrep Kit (Roche) using TruSeq DNA indexes (Illumina). Libraries were size selected at 350 – 700 bp and sequenced on the MiSeq platform as paired-end 300 bp runs at Florida State University.

#### Reference assemblies and phylogenetic analyses

Four different datasets were generated for *Pocillopora* phylogenetic analyses: a concatenated genomic loci dataset, complete mitochondrial genomes, and two single nucleotide polymorphism (SNP) datasets; one in which SNPs were removed based on proximity (unlinked SNP dataset) and one in which SNPs were not removed based on proximity (linked SNP dataset). Our concatenated genomic loci dataset comprised 57 individuals, which included the 44 *Pocillopora* sampled from the fore reef of Mo’orea, plus 11 additional *Pocillopora* individuals from Johnston et al. (2017) that were collected from Eastern Australia (*P. acuta, P. damicornis, P. verrucosa),* Hawai‘i (*P. acuta, P. damicornis, P. verrucosa, P. grandis, P. meandrina, P. ligulata)*, Clipperton Atoll (haplotype 2), and Mexico (*P. grandis*), and two outgroup individuals collected from Eastern Australia (*Seriatopora hystrix* and *Stylophora pistillata)*. ipyrad v0.9.60 (Eaton and Overcast 2020) was used to generate the concatenated genomic loci dataset by mapping to the *P. verrucosa* genome (Buitrago-López et al. 2020) with a clustering threshold of 85%, a minimum depth of eight reads to call majority rule base calls, a minimum of eight samples per locus, a maximum allowed SNP threshold of 20% per locus, and a maximum of 50% heterozygous sites per locus. The number of raw reads obtained per library ranged from 326,778 – 7,780,151. After trimming, the number of reads that mapped to the *P. verrucosa* genome (Buitrago-López et al. 2020) using ipyrad v0.9.60 (Eaton and Overcast 2020) ranged from 101,804 – 6,146,286 per library, with the number of loci in the concatenated genomic dataset ranging from 176 – 17,689 among samples, resulting in a sequence matrix consisting of 92.5% missing sites. ExaBayes v1.4.1 (Aberer et al. 2014) with default parameters was used to generate a Bayesian phylogeny of this concatenated genomic loci dataset.

To generate the dataset of complete mitochondrial genomes, all 57 libraries were mapped to the *P. grandis* mitochondrial genome, accession number EF526303 (Flot and Tillier 2007), using bwa mem v0.7.17 (Li and Durbin 2009). Consensus sequences were generated using bcftools mpileup and bcftools call (Li 2011) and converted to fasta format using Seqtk (https://github.com/lh3/seqtk). A single individual with 100% coverage of the reference was chosen per mtORF haplotype, with the exception of *P. acuta* and *P. damicornis*, for which previously sequenced complete mitochondrial genomes were used, accession numbers NC_009797 and EU_400213, respectively (Flot and Tillier 2007; Chen et al. 2008). Two outgroups, *Seriatopora caliendrum* (EF633601) and *S. hystrix* (EF633600) (Chen et al. 2008), were also included. Mitochondrial genomes were aligned in Geneious v9.1.8. A Bayesian phylogeny of this dataset was generated using BEAST v2.6.2 (Bouckaert et al. 2014) with the GTR model of evolution, a random local clock, and the birth-death model as the tree prior. The MCMC was run for 10,000,000 generations with sampling every 1,000 steps and the first 10% was removed as burn-in. A maximum likelihood phylogeny was generated using RAxML-NG (Kozlov et al. 2019) with the GTR+G model and 10,000 bootstrap replicates.

To generate an unlinked SNP dataset, trimmed reads of all libraries from step two of ipyrad v0.9.60 (Eaton and Overcast 2020) were aligned to the *P. verrucosa* genome (Buitrago-López et al. 2020) using bwa mem v0.7.17 (Li and Durbin 2009). The unlinked SNP dataset included *P. ligulata*, mtORF haplotypes 2, 3a, 3b, 3f, 3h, 8a, 10, and 11, *P. meandrina*, and *P. grandis*. SAMtools (Li et al. 2009) was used to convert sam files to sorted bam files. PCR duplicates were removed with MarkDuplicates and then variants were called with HaplotypeCaller using GATK v4.2 (McKenna et al. 2010; Van der Auwera et al. 2013). The variant call file (VCF) was subsequently filtered using VCFtools v0.1.6 (Danecek et al. 2011). For mtORFs that had more than three individuals sequenced per haplotype, only the three individuals with the least missing data were retained. *Pocillopora acuta*, *P. damicornis*, and outgroup taxa were removed from this dataset entirely due to large amounts of missing data (>80%) relative to the other libraries. The VCF was filtered to remove sites with missing data and indels, and then sites were thinned by 5,000 bp to extract unlinked SNPs. The unlinked SNP dataset for SNAPP contained 2,770 SNPs. The VCF file was converted to binary nexus format using the python script vcf2phylip v2.0 (Ortiz 2019). To generate a species tree, we used the SNAPP plugin in BEAST2 v2.6.2, which implements the multispecies coalescent model (Bouckaert et al. 2014). All taxa were treated as distinct species, *u* and *v* were calculated from the data, and a gamma distributed prior with α = 2 and β = 200 was used. The analysis ran for 4,000,000 Markov chain Monte Carlo (MCMC) generations, sampling every 1,000 steps. Convergence was assessed with Tracer v1.6 and the first 10% were removed as burn-in.

#### Discriminant Analysis of Principle Components

We also generated a dataset that retained linked SNPs, i.e., SNPs that were not thinned by proximity, for Discriminant Analysis of Principle Components (DAPC) to distinguish clusters of *Pocillopora*. We used a linked SNP file for DAPC analysis because DAPC does not use a model of evolution and therefore makes no assumptions about the underlying population genetics model, for example, those concerning linkage equilibrium or Hardy-Weinberg equilibrium. To generate the linked SNP dataset, we filtered the above VCF file (before filtering for SNAPP) using VCFtools v0.1.6 (Danecek et al. 2011) by first removing individuals from (Johnston et al. 2017) that previously sequenced on the Illumina GAIIx platform in 2013 (SS1, SD6, SD2, Pacu01, Pacu02, R17, SD1, SD4, J001, and R16). SNPs were retained if they were biallelic, had no more than 25% missing data, and had a mean sequencing depth of greater than or equal to six. In total, 7,887 SNPs were used for the DAPC analysis. Using the package adegenet v.2.1.3 in R (Jombart and Collins 2015), we used the function *find.clusters* to find the best number of *K*, which was chosen based on the lowest Bayesian Information Criterion (BIC). The number of PC axes was determined using the *optim.a.score* function. Seven PCA axes and four discriminant analysis axes were retained.

Additionally, we used nQuire (Weib et al. 2018) to investigate ploidy with the linked SNP dataset and found that a diploid model was the best fit for all *Pocillopora* libraries.

### Symbiodiniaceae analyses

#### Taxon sampling

Tissue samples from 217 *Pocillopora* colonies were collected from the fore reef of Mo’orea at LTER sites 1 – 6 in 2019. These samples included nine mtORF haplotypes (haplotype 1a, which includes *P. meandrina* and *P. grandis*, as well as 2, 3a, 3b, 3f, 3h, 8a, 10, and 11). *Pocillopora acuta* can be found on the fringing reef at Mo’orea (Rouzé et al. 2019) but was not included in our collection because we only sampled the fore reefs. The 44 *Pocillopora* samples used for phylogenetic analyses are a subset of these 217 samples. For four *Pocillopora* lineages (haplotype 10, the lineage containing *P. meandrina* and haplotype 8a, *P. grandis*, and the lineage containing haplotypes 11/2), samples were collected at 5, 10, and 20 m depths (Supplemental table 2 and (Johnston et al. 2021)).

#### DNA extraction, ITS2 library preparation, and sequencing

Genomic DNA was extracted from tissues using Chelex 100 (Bio-Rad, USA). Samples were incubated in 150 μL of 10% Chelex 100 for 60 min at 55 °C followed by 15 min at 95 °C. The supernatant was then used for PCR amplification of the ribosomal internal transcribed spacer 2 (ITS2) region using the SYM_VAR primer pairs (Hume et al. 2018). Initial PCR was performed using Phusion High-Fidelity MasterMix (ThermoFisher Scientific, USA) with an initial denaturation step at 98°C for 2 minutes, followed by 35 cycles of 98°C for 10 s, 56°C for 30 s, and 72°C for 30 s, and a final extension step at 72°C for 5 minutes. Amplified DNA was cleaned using AMPure XP beads and used as template for an index PCR in which unique combinations of Nextera XT index primers were used for each sample. Eight cycles of PCR were performed as described above with an annealing temperature of 55°C. Libraries were then cleaned, normalized, and pooled for sequencing on an Illumina MiSeq platform with 2×300 paired end reads.

Demultiplexed forward and reverse .fastq files were passed to SymPortal (Hume et al. 2019) remotely, which removed non-Symbiodiniaceae sequences and then grouped Symbiodiniaceae sequences by genera. Being a ribosomal gene, there are usually multiple ITS2 copies within a single Symbiodiniaceae cell. ITS2 sequence variation can therefore come from intragenomic variation among gene copies, in addition to intergenomic variation among Symbiodiniaceae genotypes. To parse ITS2 sequence variation into intra- and intergenomic variation, SymPortal first identifies within-sample informative intragenomic sequences, termed ‘defining intragenomic variants’ (DIVs). The repeated co-occurrence of DIVs with similar relative abundances within samples is then used to predict ITS2 type profiles, which collapse purportedly intragenomic sequence variation into taxonomic units that may represent distinct groups above, at, or below the species level (Hume et al. 2019). Since ITS2 is not a species-level marker, the psbA^ncr^ marker was also used to better resolve Symbiodiniaceae species-level diversity.

#### Cladocopium *species identification*

To complement information provided by ITS2 type profiles and increase certainty in species identification, we also used the non-coding plastid minicircle (psbA^ncr^) to distinguish between *Cladocopium* species (Moore et al. 2003) for 66 *Pocillopora* colonies in our collection. These 66 *Pocillopora* colonies encompassed the diversity of ITS2 type profiles obtained from SymPortal for which *Cladocopium* was dominant. The psbA^ncr^ region was amplified using the primers and protocol of (Moore et al. 2003), and the region was sequenced in both the forward and reverse directions. Sequences were aligned and manually checked in Geneious v.9.1.8 (Biomatters Ltd, Auckland, NZ). To add further confidence in *Cladocopium* species identification, we added 41 psbA^ncr^ sequences from (Turnham et al. 2021), which encompass a wide geographic range across the Pacific, including two *C. goreaui* sequences used as an outgroup, and three sequences from (Pinzon and Lajeunesse 2011) collected from *Pocillopora* haplotype 2 colonies from Clipperton Atoll. A Bayesian phylogeny of this larger dataset was generated using BEAST v2.6.2 (Bouckaert et al. 2014). We used the GTR model of evolution, a random local clock, and the birth-death model as the tree prior. The MCMC was run for 266,000,000 generations with sampling every 1,000 steps and the first 20% was removed as burn-in. Clades within *C. latusorum* and *C. pacificum* were identified by high posterior support (>95%) at deep monophyletic nodes, though divergence between these clades was greater in *C. latusorum* than in *C. pacificum*.

#### *Analyses of Symbiodiniaceae among* Pocillopora *species*

We used Principle Coordinate Analysis (PCoA; *cmdscale* function in the R package vegan (Oksanen et al. 2019)) with a Bray-Curtis dissimilarity measure to visualize differences in Symbiodiniaceae ITS2 sequence diversity between *Pocillopora* species or haplotype. Differences between groups were tested using Permutational Multivariate Analysis of Variance (PERMANOVA) with 99,999 permutations (*adonis* function in the R package vegan). These analyses were run to only include colonies dominated by either *C. latusorum* or *C. pacificum*.

#### *Analyses of Symbiodiniaceae within* Pocillopora *species across depth*

We also investigated whether *Pocillopora* species maintain specificity in their Symbiodiniaceae composition across depth (5, 10, and 20 m) for four *Pocillopora* species at Mo’orea. These included two species with similar relative abundances across depth (*P. grandis*, and haplotypes 11/2), and two species with different relative abundances across depth (haplotype 10, which is most prevalent at 20 m depth, and *P. meandrina* and haplotype 8a, which is most prevalent at 5 m depth) (Johnston et al. 2021). We used PCoA with a Bray-Curtis dissimilarity measure to visualize differences in Symbiodiniaceae ITS2 sequence diversity within *Pocillopora* species across depth. The vegan *pairwise.adonis2* function with 99,999 permutations was used to investigate pairwise differences across depth. As above, these analyses were run to only include colonies dominated by either *C. latusorum* or *C. pacificum*, depending on the *Pocillopora* species. Within each *Pocillopora* species, differences in ITS2 type profiles were also investigated across depth for those colonies found to host only *C. latusorum* or *C. pacificum* using Pearson’s Chi-squared Test for Count Data (function *chisq.test* function in R).

### *Cophylogenetic analysis of* Cladocopium – Pocillopora

To test for phylogenetic congruence, *Pocillopora* host and *Cladocopium* symbiont phylogenetic trees, based on psbA^ncr^, were assessed using the Procrustean Approach to Cophylogeny, PACo (R package paco (Balbuena et al. 2013)). PACo is a distance-based global-fit method that quantifies the topological congruence between two phylogenetic trees and identifies the particular associations contributing to the cophylogenetic structure. The method allows for multiple host-symbiont associations and does not require fully-resolved phylogenies. Instead of testing for random association between the host and symbiont taxa, PACo explicitly tests the dependence of the symbiont phylogeny on the host phylogeny, which assumes that symbiont speciation is driven by host speciation. PACo does this by carrying out a PCoA on the host and symbiont phylogenetic distances separately, then using Procrustes superimposition to rotate and scale the symbiont tree to fit that of the host, tests for the congruence of host – symbiont associations. The null hypothesis tested is that the algal symbiont branches are randomly associated to the coral host branches. The alternative hypothesis is that at least some part of the symbiont ordination is constrained by that of the corals and, thus, the coral-algae associations are mirrored in phylogenetic congruence. The analysis of the residual sums of squares of each topographical link between algal and host trees provides a direct measure of the importance of each specific link to the global fit and is useful for quantifying evidence for which symbiont species or genetic clades track corals more than others. Smaller residual sums of squares mean greater importance. Jackknife 95% confidence intervals also are provided. Including psbA^ncr^ sequences from multiple colonies, rather than from just one representative colony for each *Pocillopora* species, allowed us to investigate lineage diversity both within *Cladocopium* species as well as *Pocillopora* species specificity with a *Cladocopium* species or lineage.

For *Cladocopium*, we used the psbA^ncr^ Bayesian tree of 110 taxa described above, but only included symbiont – host links for the 66 taxa we sequenced the psbA^ncr^ region for in the PACo analysis. For *Pocillopora*, a Bayesian phylogenetic tree was generated using ExaBayes v1.4.1 (Aberer et al. 2014) with default parameters by selecting a single representative for *P. verrucosa*, *P. meandrina*, *P. grandis*, and haplotypes 2, 8a, and 11 with the highest coverage from the concatenated genomic loci dataset generated by ipyrad. Phylogenetic trees were imported into R using the “ape” package (Paradis et al. 2004).

## Results

### Pocillopora *phylogenetic analyses*

Analyses using two approaches to delineate *Pocillopora* species (DAPC analysis using 7,887 linked SNPs and SNAPP analysis using 2,770 unlinked SNPs) supported five species on the fore reef at Mo’orea. Furthermore, *P. ligulata* from Hawai’i was distinct from any species at Mo’orea and does not occur there (Figures 1 and 2). The five species of *Pocillopora* recovered from the fore reef at Mo’orea were: 1) haplotypes 11/2, 2) *P. meandrina* and haplotype 8a, 3) *P. verrucosa* (haplotypes 3a, 3b, 3f, and 3h), 4) Haplotype 10, and 5) *P. grandis*. Note that *P. acuta* also occurs at Mo’orea, but only on the fringing reef (Rouzé et al. 2019). In total, there are six species of *Pocillopora* at Mo’orea.

**Figure 1.**
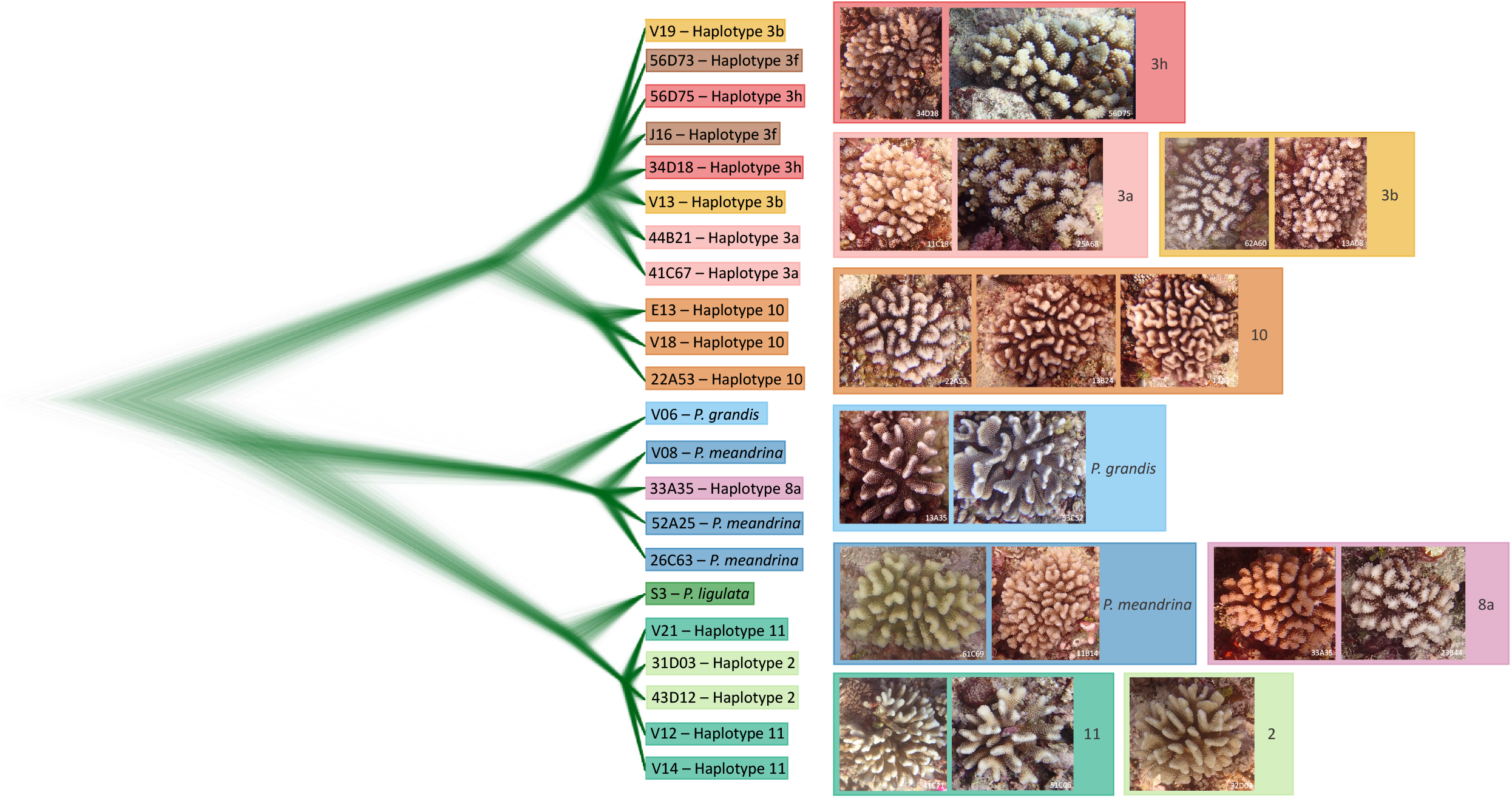
*Pocillopora* species tree. The cloudogram distribution from SNAPP showing the Bayesian analysis of 2,770 nuclear, unlinked, bi-allelic SNPs.

**Figure 2.**
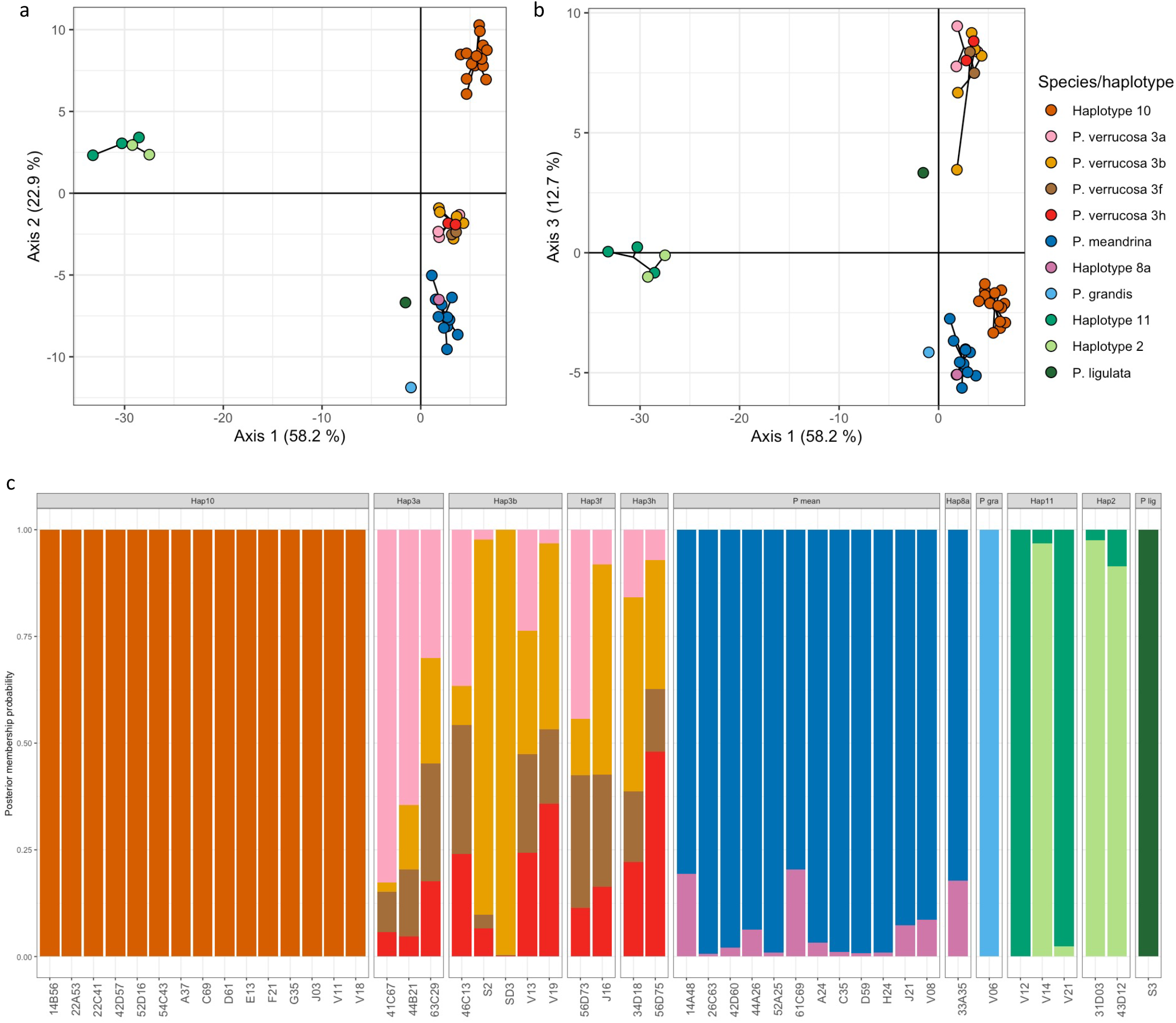
Discriminant Analysis of Principle Components (DAPC) plots for *Pocillopora* using 7,887 nuclear bi-allelic SNPs that were not filtered for linkage showing genetic clusters in **a)** and **b)**, and the posterior membership probability of individuals to these DAPC clusters in **c)**.

The first three of these species each formed distinct clusters indicating ongoing or recent gene flow among the mtORF haplotype lineages within each cluster (Figures 1 and 2). Within *P. verrucosa*, there was no consistent clustering of haplotypes 3a, 3b, 3f, and 3h (Figures 1; 2; 3a, b). In all datasets (genomic, complete mitochondrial genomes, DAPC and SNAPP), haplotype 10 was consistently recovered as distinct from *P. verrucosa* (Figures 1; 2; 3a, b), indicating that it is a genetically distinct sister species to *P. verrucosa*.

In the mitochondrial dataset, haplotype 2 was recovered as basal to the clade containing *P. ligulata* and haplotype 11 (Figure 3b), but in the genomic (Figure 3a) and SNAPP (Figure 1) datasets *P. ligulata* was recovered as basal to haplotypes 11 and 2. In the genomic dataset, haplotypes 11 and haplotype 2 were recovered as separate monophyletic lineages (Figure 3a), but in the SNAPP dataset this pattern was not supported (Figure 1). The DAPC analysis indicated that one haplotype 11 individual hosted greater posterior membership probability to haplotype 2 than haplotype 11 (Figure 2).

**Figure 3.**
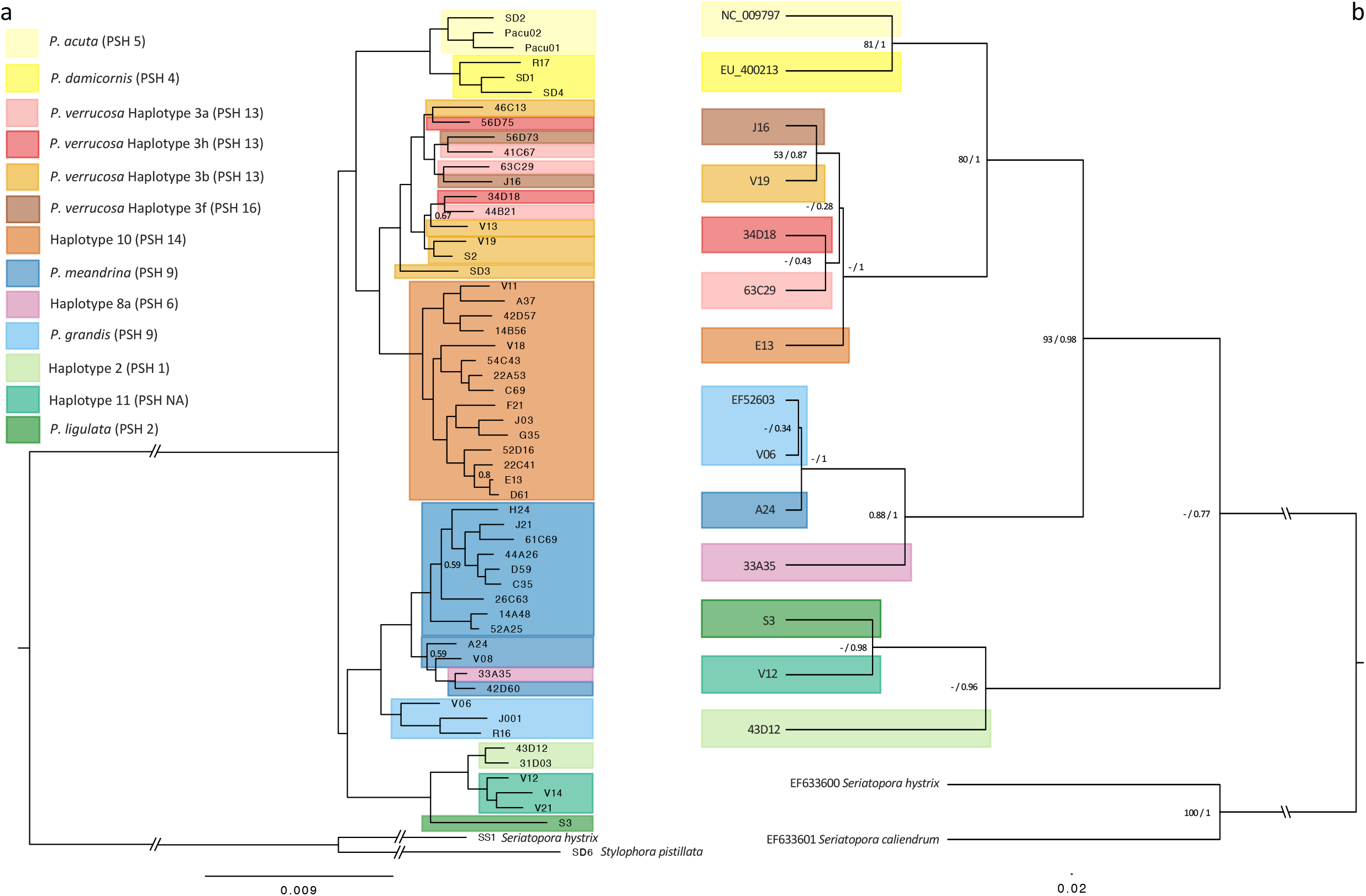
*Pocillopora* genomic and mitochondrial phylogenies. **a)** Nuclear genomic Bayesian phylogeny of *Pocillopora* with outgroups *Seriatopora* and *Stylophora*. All nodes are >95% CI unless otherwise indicated. **b)** Maximum and Bayesian phylogenetic analysis of complete mitochondrial genomes. Mitochondrial haplotype identification follows (Pinzón et al. 2013) and (Forsman et al. 2013). For ease of comparison among studies using different naming, the Primary Species Hypothesis (PSH) presented in (Gélin et al. 2017) is added in parentheses.

In both the genomic and SNAPP datasets, *P. grandis* was recovered as basal to the clade that contains *P. meandrina* and haplotype 8a, but in the mitochondrial dataset haplotype 8a was basal to *P. meandrina* and *P. grandis* (Figure 3b). Haplotype 8a was not differentiated from *P. meandrina* in the genomic, SNAPP, or DAPC datasets.

### Pocillopora – *Symbiodiniaceae specificity*

The majority (215 of 217) of *Pocillopora* colonies sampled from Mo’orea hosted Symbiodiniaceae from the genus *Cladocopium* (Figure 4). Four out of 41 (10%) *P. grandis* colonies hosted *Symbiodinium* spp. and one hosted *Durusdinium* spp. (2%). Five out of 26 (19%) haplotype 11 and haplotype 2 colonies hosted *Durusdinium glynnii*. The undescribed *Cladocopoium* species identified with the C116 DIV was hosted by one *P. verrucosa* colony, two *P. meandrina* colonies, one *P. grandis* colony, and one haplotype 11 colony (Figure 4).

**Figure 4.**
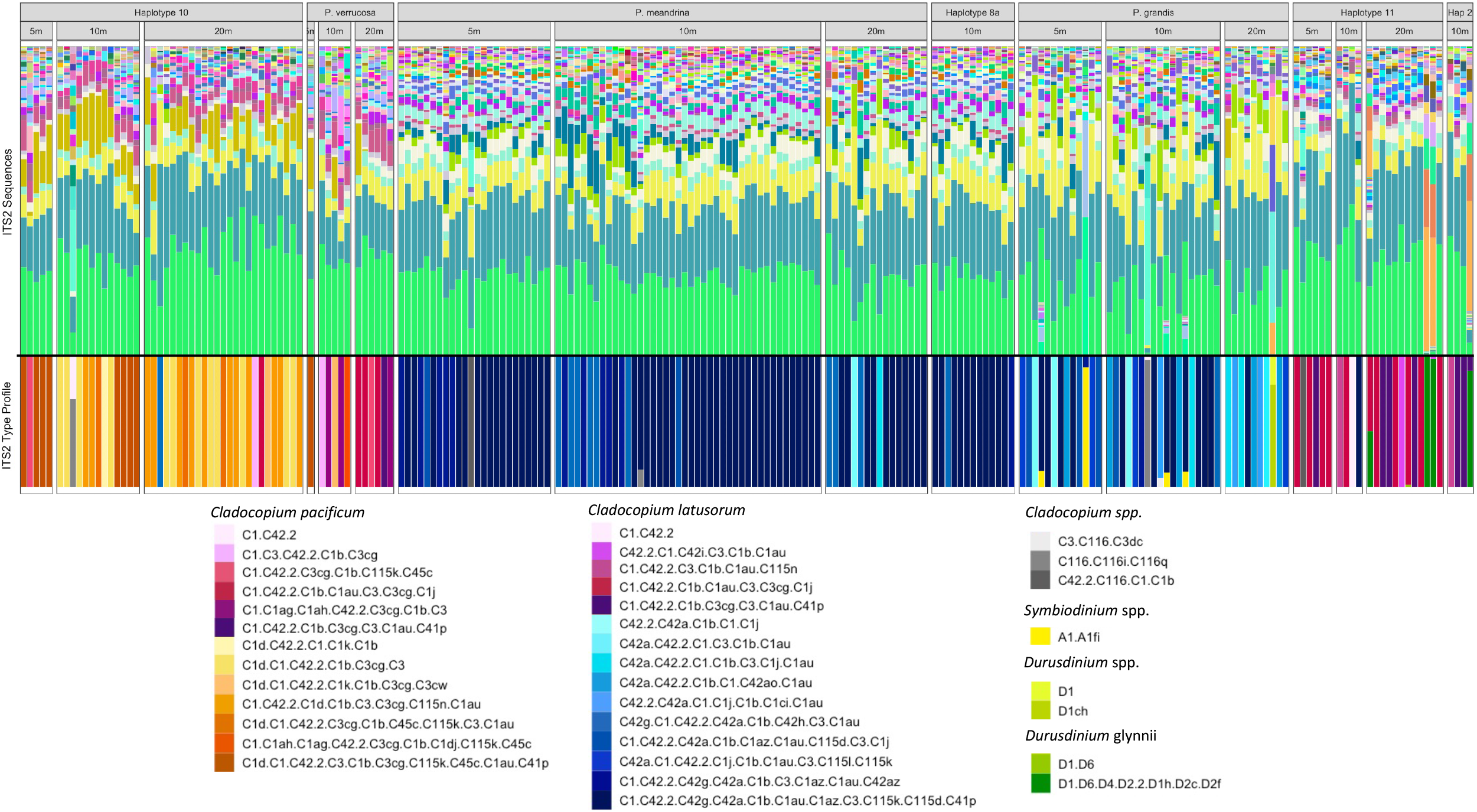
Diversity of ITS2 sequences and ITS2 type profiles within each colony of *Pocillopora* species or haplotype. Each column represents a colony and colonies are binned by depth sampled within each species or haplotype. ITS2 sequence color key found in Supplemental Figure 3.

Phylogenetic analysis of the *Cladocopium* psbA^ncr^ region indicated that the majority of symbionts hosted by *Pocillopora* were either *C. latusorum* or *C. pacificum* (Figure 5a). Sequences from colonies hosting C116 DIVs could not be aligned with those from *C. latusorum* and *C. pacificum*, and therefore were not included in the phylogenetic analysis. *Cladocopium latusorum* was hosted by *P. meandrina* and haplotype 8a, *P. grandis*, and haplotypes 11/2, and one haplotype 10 colony. *Cladocopium pacificum* was hosted exclusively by haplotype 10 and *P. verrucosa*. Largely, and following (Turnham et al. 2021), we found that *C. latusorum* could be identified by the C42a DIV in their ITS2 type profile, and that *C. pacificum* could be identified by the C1d DIV in their profile (Figures 4 and 5a). However, there were also some ITS2 type profiles from haplotype 10, *P. verrucosa*, and haplotypes 11/2 colonies that could not be assigned to a *Cladocopium* species solely based on their ITS2 type profiles because they did not contain C42a or C1d. These ambiguous ITS2 type profiles were C1.C42.2, C1.C3.C42.2.C1b.C3cg, C42.2.C1.C42i.C3.C1b.C1au, C1.c42.2.C3cg.C1b.C115k.C45c, C1.C42.2.C3.C1b.C1au.C115n, C1.C42.2.C1b.C1au.C3.C3cg.C1j, C1.C1ag.C1ah.C42.2.C3cg.C1b.C3, and C1.C42.2.C1b.C3cg.C3.C1au.C41p. These profiles, however, could be assigned to either *C. latusorum* or *C. pacificum* using the pbsA^ncr^ marker. Additionally, by quantifying the presence of ITS2 DIVs in each colony, we found that there are unique DIV associations with *Pocillopora* species (Supplemental figure 1) and ITS2 type profiles (Supplemental figure 2) that were not included in the ITS2 type profiles predicted by SymPortal.

**Figure 5.**
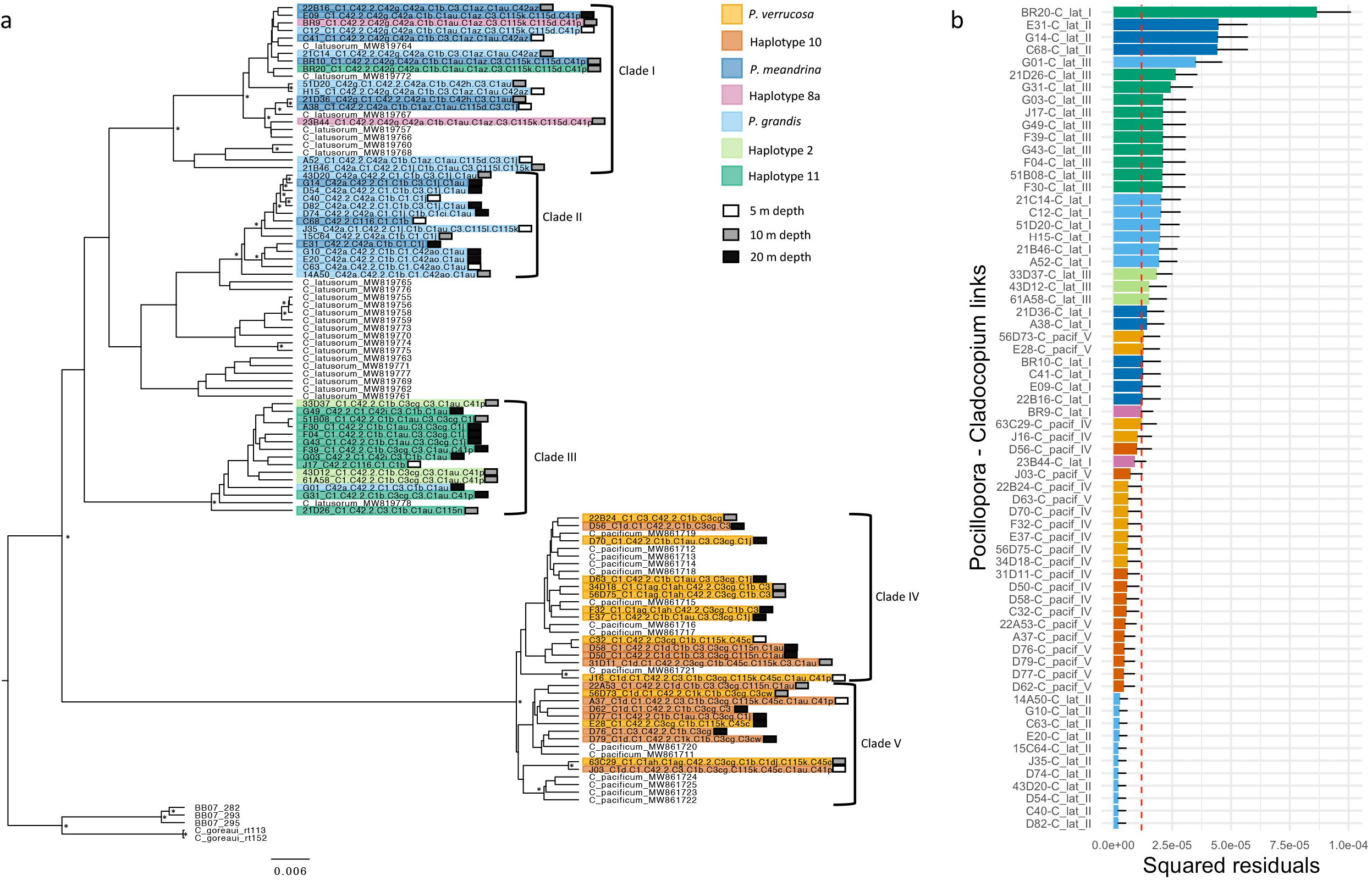
*Cladocopium* psbA Bayesian tree and PACo Procrustean analysis of host symbiont links. **a)** *Cladocopium* psbA Bayesian tree that includes 66 samples from this study (those identified in color by *Pocillopora* species/haplotype). Depth of collection is indicated to the right of taxa labels. Taxa labeled as BB07 are from Pinzon and LaJeunesse 2011 and all other taxa are from Turnham et al. 2021. **b)** Contributions of individual *Pocillopora* host – *Cladocopium* symbiont links to the Procrustean fit. Jackknifed squared residuals (bars) and upper 95% confidence intervals (error bars) resulting from applying PACo to symbiont distances. The *Pocillopora* species/haplotype each psbA sequence was extracted from are indicated by the bar color. The median squared residual value is shown (red dashed line).

#### *Symbiodiniaceae composition among* Pocillopora *species*

ITS2 sequence diversity differed significantly between most *Pocillopora* species and haplotypes (Figure 6a, Table 1). The greatest difference in ITS2 sequence diversity was between the *Pocillopora* lineage that contains *P. meandrina*, haplotype 8a, and *P. grandis*, and the other *Pocillopora* species and haplotypes. Symbiodiniaceae ITS2 diversity did not differ between *P. meandrina* and haplotype 8a (*p* = 0.371), or between haplotypes 11 and 2 (*p* = 0.888), providing further support for hybridization or introgression between these *Pocillopora* groups.

**Figure 6.**
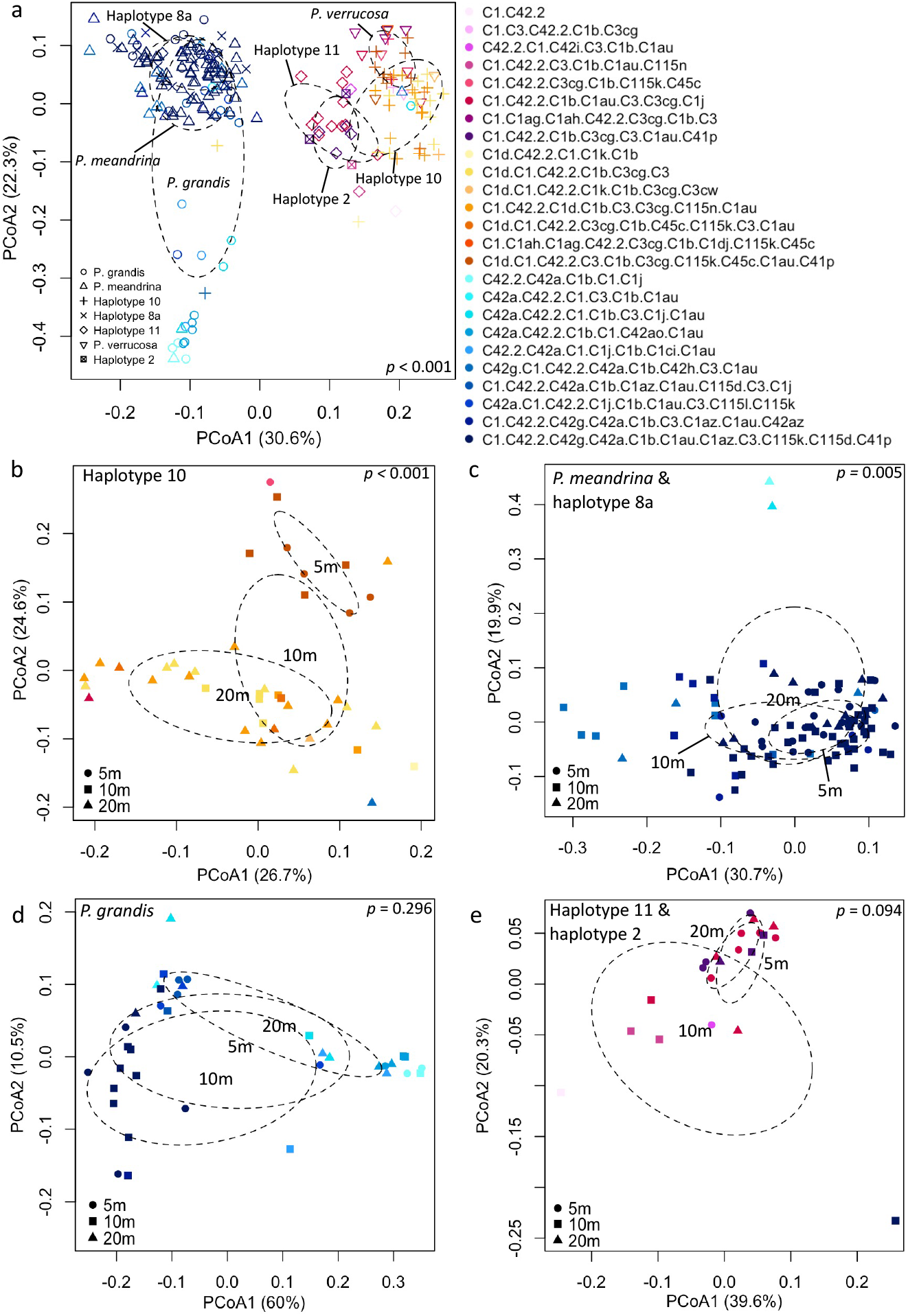
Principle Coordinates Analyses of Symbiodiniaceae ITS2 sequence diversity for **a)** *Pocillopora* species and mtORF haplotypes sampled from the fore reefs of Mo’orea, and then across depth for **b)** haplotype 10, **c)** *P. meandrina* and haplotype 8a, **d)** *P. grandis*, and **e)** haplotypes 11/2. Symbols are colored by ITS2 type profile and PERMAMOVA results are presented in the top or bottom right corner of each plot.

**Table 1.** PERMANOVA results for ITS2 sequence diversity for *Pocillopora* species and haplotypes, and for *Pocillopora* species and haplotype across depth, when only colonies with ITS2 type profiles from *C. latusorum* and *C. pacificum* were included. *Pocillopora verrucosa* was not included in depth analyses because of low sample size. Df - degrees of freedom; Sum Sq - sum of squares; Pseudo-F - F value by permutation, boldface indicates statistical significance with P < 0.05, P-values based on 99,999 permutations.

|  | Df | Sum Sq | Pseudo-F | R<sup>2</sup> | P |
| --- | --- | --- | --- | --- | --- |
| *Pocillopora* species/haplotype | 6 | 1.294 | 18.469 | 0.356 | **< 0.001** |
| Residuals | 201 | 2.346 |  | 0.644 |  |
| Haplotype 10 by depth | 2 | 0.068 | 3.247 | 0.143 | **< 0.001** |
| Residuals | 39 | 0.408 |  | 0.857 |  |
| *P. meandrina* and haplotype 8a by depth | 2 | 0.039 | 2.426 | 0.051 | **0.005** |
| Residuals | 91 | 0.732 |  | 0.949 |  |
| *P. grandis* by depth | 2 | 0.045 | 1.160 | 0.062 | 0.296 |
| Residuals | 35 | 0.676 |  | 0.938 |  |
| Haplotype 11 and haplotype 2 by depth | 2 | 0.018 | 1.535 | 0.146 | 0.095 |
| Residuals | 18 | 0.106 |  | 0.854 |  |

#### *Symbiodiniaceae composition within* Pocillopora *species across depth*

Within *Pocillopora* species, ITS2 sequence diversity differed across depth for some *Pocillopora* species but not others (Figure 6b-e, Tables 1 and 2). Pairwise comparisons of ITS2 sequence diversity were significant within *C. pacificum* hosted by haplotype 10 between 5 – 10 and 5 – 20 m depths (Figure 6b, Table 2). Additionally, Pearson’s Chi-squared Test indicated that ITS2 type profiles within *C. pacificum* also differed significantly across depth (*X*^2^ (20) = 36.917; *p* = 0.012). Within *P. meandrina* and haplotype 8a, pairwise comparisons of ITS2 sequence diversity within *C. latusorum* were significant between 5 – 20 and 10 – 20 m depths (Figure 6b, Table 2). However, ITS2 type profiles did not differ significantly across depth. ITS2 sequence diversity did not differ across depth in *P. grandis* or haplotypes 11/2 (Table 2).

**Table 2.** Pairwise comparisons of ITS2 Sequence diversity for *Pocillopora* species and haplotypes across depth. ITS2 sequence diversity analyses include sequences when only colonies with ITS2 type profiles from *C. latusorum* and *C. pacificum* were included. Df - degrees of freedom; Sum Sq - sum of squares; Pseudo-F - F value by permutation, boldface indicates statistical significance with P < 0.05, P-values based on 99,999 permutations.

|  | 5 - 10m |  |  |  |  | 5 - 20m |  |  |  |  | 10 - 20m |  |  |  |  |
| --- | --- | --- | --- | --- | --- | --- | --- | --- | --- | --- | --- | --- | --- | --- | --- |
|  | Df | Sum Sq | Pseudo-F | R<sup>2</sup> | P | Df | Sum Sq | Pseudo-F | R<sup>2</sup> | P | Df | Sum Sq | Pseudo-F | R<sup>2</sup> | P |
| **Haplotype 10** | 1 | 0.025 | 2.872 | 0.161 | **0.032** | 1 | 0.056 | 5.061 | 0.153 | **0.001** | 1 | 0.020 | 1.888 | 0.051 | 0.069 |
| Residuals | 15 | 0.131 |  | 0.839 |  | 28 | 0.309 |  | 0.847 |  | 35 | 0.376 |  | 0.949 |  |
| ***P. meandrina* and haplotype 8a** | 1 | 0.015 | 2.078 | 0.027 | 0.054 | 1 | 0.018 | 2.310 | 0.059 | **0.028** | 1 | 0.026 | 2.771 | 0.039 | **0.015** |
| Residuals | 76 | 0.533 |  | 0.973 |  | 37 | 0.295 |  | 0.941 |  | 69 | 0.636 |  | 0.961 |  |
| ***P. grandis*** | 1 | 0.012 | 0.592 | 0.021 | 0.63 | 1 | 0.016 | 0.934 | 0.047 | 0.357 | 1 | 0.039 | 1.975 | 0.076 | 0.106 |
| Residuals | 27 | 0.548 |  | 0.979 |  | 19 | 0.331 |  | 0.953 |  | 24 | 0.474 |  | 0.924 |  |
| **Haplotype 11 and haplotype 2** | 1 | 0.012 | 1.380 | 0.121 | 0.192 | 1 | 0.003 | 1.090 | 0.083 | 0.347 | 1 | 0.012 | 1.841 | 0.116 | 0.082 |
| Residuals | 10 | 0.085 |  | 0.879 |  | 12 | 0.034 |  | 0.917 |  | 14 | 0.093 |  | 0.884 |  |

### *Cophylogenetic analysis of* Cladocopium – Pocillopora

We found strong dependence of the *Cladocopium* phylogeny on the *Pocillopora* phylogeny (Figure 7). The test for the extent to which the coral – algal associations are mirrored in phylogenetic congruence yielded a global residual sum of squares (*m*^2^) of 0.0009 with a permutational p-value of *p* < 0.0001, indicating that the *Cladocopium* phylogeny is not randomly associated with the *Pocillopora* phylogeny. We also identified three clades within *C. latusorum* and two clades within *C. pacificum*.

**Figure 7.**
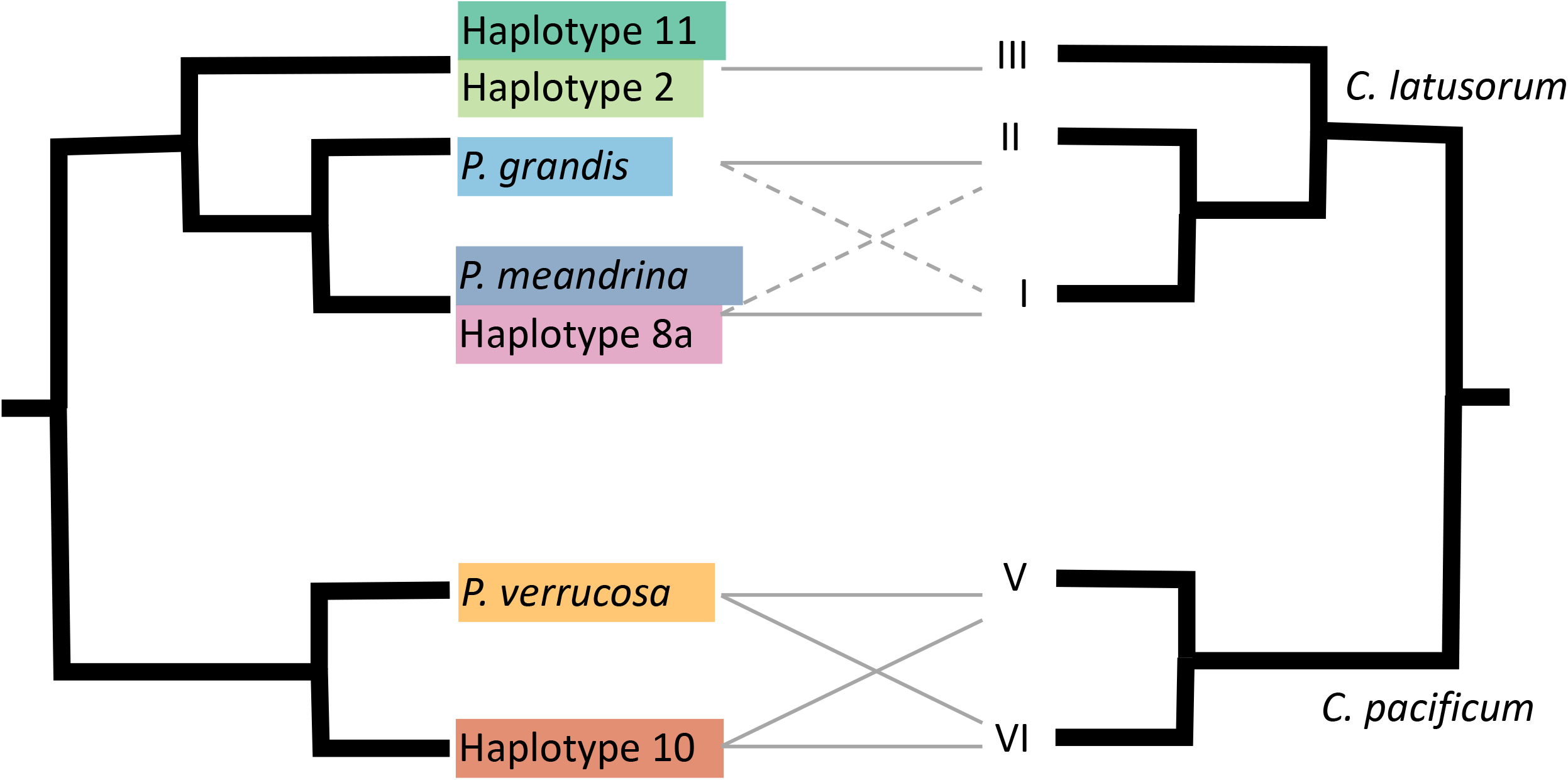
Schematic summary of *Pocillopora* host – *Cladocopium* symbiont cophylogeny, based off analyses shown in Figure 6. Dashed lines indicate minor links between host and symbiont clades reflecting greater flexibility.

The specific coral – algal associations that contributed the most (i.e., smallest squared residuals) to the global phylogenetic congruence were the dependence of *C. latusorum* clade II on *P. grandis*, and the dependence of *C. pacificum* clades IV and V on *Pocillopora* sister taxa haplotype 10 and *P. verrucosa*. The weakest associations contributing to the overall phylogeny were the dependence of *C. latusorum* clade III and its dependence on haplotype 11 (Figure 5b). Although there was high specificity between haplotypes 11/2 and *C. latusorum* clade III, the contribution of these links was low because of the low branch support within this clade.

## Discussion

Identifying the dependence of symbiont lineages on their host lineages is important for understanding the evolutionary history and ecological responses of the holobiont (Corbin et al. 2017). Despite the obligate symbiotic relationships between photosynthetic algae (Symbiodiniaceae) and scleractinian coral hosts being a classic example of a holobiont, explicit tests of phylogenetic congruence have been absent. Furthermore, tests of cophylogeny between corals and their algal symbionts have been hampered in the past because both corals and algae contain genetically unresolved and morphologically-cryptic species. Here, we used multiple genomic datasets from samples collected across multiple depths and sites at Mo’orea, French Polynesia, to resolve previously well-studied mitochondrial lineages into species and found strong evidence of cophylogeny between *Pocillopora* species and their algal symbionts. Our analyses supported the presence of five co-occurring *Pocillopora* species on the fore reef of Mo’orea that largely hosted two species of host-specialized algae (*Cladocopium latusorum* and *C. pacificum). Pocillopora acuta* can also be found at Mo’orea but is only found on the fringing reef, not on the fore reef, and largely hosts *Durusdinium* in this environment (Rouzé et al. 2019; Strader et al. 2022) and was therefore not included in our analyses. In general, the *Cladocopium* phylogeny, including multiple clades within species, mirrored the *Pocillopora* phylogeny (Figure 7). Certain *Cladocopium* species and lineages within species showed stronger dependence with specific *Pocillopora* species than others. Such cophylogeny is likely a consequence of the life history of broadcast-spawning *Pocillopora*, which, unlike many broadcast spawning corals from other genera, transmit algal symbionts to the next generation in the eggs (Hirose et al. 2000; Schmidt-Roach et al. 2012). However, we also found that some *Pocillopora* colonies were dominated by symbiont taxa from *Symbiodinium* and *Durusdinium*. Taxa from *Symbiodinium* and *Durusdinium* also occur in other host taxa ranging from jellies (Sachs and Wilcox 2006; Mammone et al. 2021) to clams (DeBoer et al. 2012; Poo et al. 2021), and may have a free-living stage (Pochon et al. 2014; Fujise et al. 2021). The presence of symbiont taxa from *Symbiodinium* and *Durusdinium* suggests that there may also be some degree of horizontal transmission in broadcast-spawning *Pocillopora*. While speciation of *C. latusorum* and *C. pacificum* appears to be driven by niche diversification provided by speciation in *Pocillopora* (Turnham et al. 2021), we are now in a position to understand more about certain associations of symbiont clades within more *Pocillopora* species. Furthermore, *Pocillopora* still harbors generalist symbionts that may also be transmitted vertically but could be acquired horizontally as well.

Recently, (Turnham et al. 2021) described two new species of Symbiodiniaceae in the genus *Cladocopium* and found that they co-diversified with different *Pocillopora* lineages, likely as a result of maternal vertical transmission of symbionts. They found that *C. latusorum* co-diversified with the lineage that contains *P. meandrina*, haplotype 8a, and *P. grandis*, and that *C. pacificum* co-diversified with *P. verrucosa*, and that this speciation event in *Cladocopium* coincides with the diversification of these extant *Pocillopora* lineages approximately 3MYA. Here, we provide additional symbiont associations for several *Pocillopora* species and provide a formal test of cophylogeny. Specifically, we found that, in addition to *P. meandrina*, haplotype 8a, and *P. grandis*, haplotypes 11/2 also host *C. latusorum*. In addition to *P. verrucosa*, haplotype 10 also hosts *C. pacificum*. Furthermore, we show that there are unique associations between *Pocillopora* species and different lineages within *C. latusorum* and *C. pacificum*, providing further evidence for the co-speciation hypothesis.

In the past, cophylogeny between corals and their algal symbionts was expected to be rare or absent (Van Oppen and Medina 2020) because many coral species exhibit horizontal transmission (Baird et al. 2009), many algal symbiont species exhibit flexibility to associate with different hosts (LaJeunesse et al. 2018), and even coral species with vertical transmission can have some degree of horizontal acquisition of symbionts (Quigley et al. 2018, 2019). Other coral species, including members from *Porites* and *Montipora*, also exhibit vertical transmission and harbor host-specialist species of *Cladocopium* (Lajeunesse et al. 2004; LaJeunesse and Thornhill 2011; Forsman et al. 2020; Hoadley et al. 2021), but the extent of cophylogeny within these groups remains to be tested. Members of the genus *Cladocopium* associate with a broad diversity of hosts in addition to corals, including other cnidarians, clams, ciliates, flatworms, foraminifera, and sponges, but can be highly host specialized (LaJeunesse et al. 2018). Therefore, an important next step to test the links between vertical transmission and co-speciation is to compare estimates of cophylogeny among more groups of corals and compare groups with vertical versus horizontal transmission.

The functional differences, if any, between *C. latusorum* and *C. pacificum* in their effects on the coral host are not clear. However, the high specificity and evidence for shared evolutionary history still generates hypotheses for the degree to which symbioses in this group promote or restrict adaptation to climate change (Compant et al. 2010). For example, the ability to switch or shuffle symbionts in response to environmental change may be lower in coral species with high symbiont specificity. To test this, it will be informative to know the extent to which *Pocillopora* species hosting *C. latusorum* or *C. pacificum* differ in their ability to switch or shuffle symbionts. In our dataset, only *Pocillopora* species that typically host *C. latusorum* showed the ability to host taxa from *Symbiodinium* and *Durusdinium*. In fact, only those with similar relative abundances between 5m and 20m (*P. grandis*, and haplotype 11 and 2 (Johnston et al. 2021)) hosted symbionts from genera other than *Cladocopium*. For example, *Durusdinium glynnii*, a heat-tolerant host generalist (LaJeunesse et al. 2010), was detected in five haplotype 11/2 colonies. *D. glynni* forms stable, long-term associations with hosts that are often found in highly variable environments (LaJeunesse et al. 2010; McGinley et al. 2012; Sawall et al. 2014; Innis et al. 2018; Rouzé et al. 2019). Interestingly, haplotypes 11/2 experienced the greatest bleaching mortality following the bleaching event at Mo’orea in 2019 (Burgess et al. 2021), thus the association with *D. glynnii* in the survivors sampled here could reflect the outcome of selective mortality or symbiont shuffling following this event. Furthermore, the decreased thermal tolerance of haplotypes 11/2 may relate to their unique association with *C. latusorum* Clade III (Figure 5), reduced fitness from introgression among hosts (Muhlfeld et al. 2009; Kim et al. 2018), or the interplay between both (Miller et al. 2010). Similarly, four *P. grandis* colonies contained 12-91% *Symbiodinium* spp. This genus contains both free-living and obligate symbionts generalist species found in many tropical environments and in a diversity of hosts such as jellyfish, giant clams, and other coral genera (LaJeunesse 2017). Some taxa are highly infectious and when attracted to certain hosts, can swarm in high densities next to these available hosts (Yamashita et al. 2014). In the sea anemone, *Exaiptasia pallida*, colonization by *S. microadriaticum* was found to be akin to a form of parasitism in which there is no apparent benefit to the host (Gabay et al. 2018). It is interesting that *Symbiodinium* was only detected in *P. grandis*. In the Red Sea, *P. verrucosa*, increasingly hosts *Symbiodinium* with decreasing latitude, indicating that there may be some benefit to this symbiosis with increasing sea water temperature (Sawall et al. 2014). But whether the *Symbiodinium* taxa hosted in *P. grandis* at Mo’orea is the same as that hosted in the Red Sea is not known. Additionally, whether this symbiosis with *P. grandis* is long-lasting or brief is not known, nor whether this *Symbiodinium* spp. is more attracted to *P. grandis* than other *Pocillopora* species at Mo’orea. Alternatively, the *Cladocopium* lineages hosted by *P. grandis* may be poorer competitors with this *Symbiodinium* spp. than those hosted by other *Pocillopora*, or the immune system of *P. grandis* may be less able to detect this *Symbiodinium* spp.

A higher rate of sexual reproduction within a *Cladocopium* species or clade could lead to greater potential for adaptation to different environments. Recently, cytological evidence of reproduction of *C. latusorum* in *P. meandrina* was found to be of mixed mode; mainly asexual with sexual reproduction occurring in < 1-5% of cells *in hospite* (Figueroa et al. 2021). While there is some evidence that sexual reproduction in free-living Symbiodiniaceae may be triggered by stress, such as nutrient deficiency (Pfiester 1989), the triggers of sexual reproduction in symbiotic Symbiodiniaceae, such as *C. latusorum* (Fujise et al. 2021), are not known. These triggers may be both biotic and abiotic, differ by *Cladocopium* species or lineage, and/or be controlled by the host. The depth differences we detected in the symbiont ITS2 sequence diversity and type profiles within *Pocillopora* haplotype 10 may indicate the potential for adaptive divergence in *C. pacificum*, as hypothesized in other coral species (Bongaerts et al. 2011, 2013).

By genetically identifying species of both *Pocillopora* and Symbiodiniaceae, our analysis overcomes a major hurdle that has prevented tests of coevolution and co-speciation in corals and their symbiotic algae. For example, we can also confirm that haplotype 10 (Forsman et al. 2013) is a distinct *Pocillopora* species that warrants formal identification. Although haplotype 10 hosted the same algal species (*C. pacificum*) as its sister species *P. verrucosa* (haplotypes 3a, 3b, 3f, and 3h), it tended to host a different psbA^ncr^ clade and ITS2 type profile. Haplotype 10 and *P. verrucosa* are both more common at 20m than 5m, though haplotype 10 is far more abundant than *P. verrucosa* at all depths at Mo’orea (Johnston et al. 2021). Furthermore, geographic sampling of *Pocillopora* to date indicates that haplotype 10 may be endemic to French Polynesia while *P. verrucosa* is widely distributed from the Tropical Eastern Pacific to the Red Sea and Arabian Gulf (Forsman et al. 2013; Mayfield et al. 2015; Gélin et al. 2017). As a result, we hypothesize that haplotype 10 diverged in sympatry from *P. verrucosa*, and that this is reflected in some of the genetic divergence in *C. pacificum* lineages among these two *Pocillopora* species.

*Pocillopora* haplotypes showing evidence for introgression and similar ecology also exhibited similar *Cladocopium* composition. In contrast to (Johnston et al. 2017), in which colonies were sampled from across the Pacific and where evidence of hybridization was found only between the youngest *Pocillopora* species (the brooders *P. acuta* and *P. damicornis*), here we found evidence of introgression in multiple branches of the phylogeny in co-occurring *Pocillopora* corals from Mo’orea, French Polynesia. Genomic data indicated fewer species than mitochondrial data. Because mitochondrial evolution proceeds more slowly than nuclear evolution in anthozoans (Shearer et al. 2002), our genomic analysis suggests relatively recent introgression between *P. meandrina* (mitochondrial haplotype 1a) and haplotype 8a, and between haplotypes 11/2, even though these haplotypes resolved as reciprocally monophyletic in our genomic analysis. Our finding of introgression between *P. meandrina* (mitochondrial haplotype 1a) and haplotype 8a also contrasts with Gélin et al. (2017) who designated haplotype 1a and haplotype 8a as distinct Primary Species Hypotheses (PSH) using mostly mitochondrial markers (Figure 3). Similarly, we found no evidence that haplotype 3f (PSH 16) is a distinct species from haplotypes 3a, 3b, 3h (PSH 13) using nuclear and mitochondrial genomes (cf (Gélin et al. 2017)). Furthermore, despite a lack of divergence in the mitochondrial genome in *P. meandrina* and *P. grandis*, these species are distinct, confirming previous findings (Johnston et al. 2017). Similarly, mitochondrial data suggests that haplotype 2 is ancestral to haplotype 11 and *P. ligulata*. Haplotype 2 is geographically widespread, found from Clipperton Atoll in the Tropical Eastern Pacific to Madagascar in the western Indian Ocean, but is relatively rare throughout this range (Pinzón et al. 2013; Marti-Puig et al. 2014; Gélin et al. 2017) and at Mo’orea (Burgess et al. 2021). In contrast, Haplotype 11 has, to date, only been sampled from French Polynesia (Forsman et al. 2013; Burgess et al. 2021; Johnston et al. 2021). Given the basal placement of haplotype 2 in the mitochondrial phylogeny, and their respective geographic distributions, we hypothesize that haplotype 11 and *P. ligulata* diverged from haplotype 2 in isolation. However, the nuclear introgression observed between haplotypes 11/2, and the lack of differentiation in symbionts hosted, suggest that there is some gene flow between these haplotypes at Mo’orea. At Clipperton Atoll, haplotype 2 hosts a yet to be described species that is different from that hosted in Mo’orea, which is neither *C. latusorum* nor *C. pacificum*, that is closely related to *C. goreaui* (Figure 5a; taxa beginning with BB07) (Pinzon and Lajeunesse 2011). Whether there are fitness impacts when interspecific crosses occur between *Pocillopora* that host different algal species, such as the failure to transmit the algal symbiont or reduced symbiont growth if transmitted, is not known. Given the emerging evidence, the mosaic of *Pocillopora* species distributions, and thus respective interactions, is likely evolutionarily complex across time and space, giving rise to lineages and species following a pattern of reticulate evolution, as has been hypothesized for many other coral species (Veron and Stafford-Smith 2000; Diekmann et al. 2001; Arrigoni et al. 2016).

## Acknowledgements

This work was funded by a National Science Foundation (NSF) grant to S.C Burgess (OCE 1829867). We thank J. Powell and C. Peters for invaluable assistance in the field, C. Peters and the Florida State University Dive Program for facilitating field work on SCUBA, the staff of the UC Berkeley Richard B. Gump South Pacific Research Station for facilitating our research. The authors declare no conflicts of interest. Research was completed under permits issued by the French Polynesian Government (Délégation à la Recherche), the Haut-Commissariat de la République en Polynésie Française (DTRT) (Protocole d’Accueil 2019), and the U.S. Fish and Wildlife. We also thank the Core Facility in the Department of Biological Science, Florida State University, for library generation and sequencing.

## Data availability

All data and R code used to create the plots can be found at https://github.com/scottcburgess/Cophylogeny-Pocillopora-Symbiodiniaceae.

## Notes

### Competing Interest Statement

The authors have declared no competing interest.

