## Supplementary material for "Cophylogeny and specificity between cryptic coral species (*Pocillopora* spp.) at Mo’orea and their symbionts (Symbiodiniaceae)": Supplememental material

Materials and Methods:

*Phylogenetic analysis and heatmap of ITS2 DIVs*: ITS2 sequences of the 94 ITS2 DIVs that SymPortal identified for ITS2 type profile generation were obtained from the SymPortal output and were aligned in Geneious v9.1.8. A Bayesian analysis of these DIVs was carried out in BEAST2 v2.6.2 (Bouckaert et al. 2014) using the GTR model of evolution, a random local clock, and the birth-death model as the tree prior. The MCMC was run for 10,000,000 generations with sampling every 1,000 steps and the first 10% was removed as burn-in. We converted colony ITS2 relative proportion to presence-absence data and then used R v3.6.2 (R Core Team, 2019) to generate a heatmap of the 94 DIVs for each *Pocillopora* species or haplotype and for each ITS2 type profile.

**Supplemental table 1.** Collection location of *Pocillopora* samples from Mo’orea, French Polynesia, used in phylogenetics analyses.

**Supplemental table 2.** Collection location of *Pocillopora* samples from Mo’orea, French Polynesia, used in Symbiodiniaceae ITS2 analyses.

**Supplemental figure 1.** Bayesian analysis and heatmap of the 94 Symbiodiniaceae ITS2 Defining Intragenomic Variants (DIVs) that SymPortal (Hume et al. 2019) used to construct ITS2 type profiles. Nodes with >95% CI are identified with an asterisk and DIVs that have been identified to species are highlighted. The heatmap shows the percentage of colonies within a given *Pocillopora* species or haplotype that were found to host each DIV.

**Supplemental figure 2.** Heatmap of the 94 Symbiodiniaceae ITS2 Defining Intragenomic Variants (DIVs) within each ITS2 type profile.

**Supplemental figure 3.** Color key for ITS2 sequences found in *Pocillopora* as shown in Figure 4.
