## Supplementary figures and images for "Cophylogeny and specificity between cryptic coral species (*Pocillopora* spp.) at Mo’orea and their symbionts (Symbiodiniaceae)"

### Supplemental Figure 1

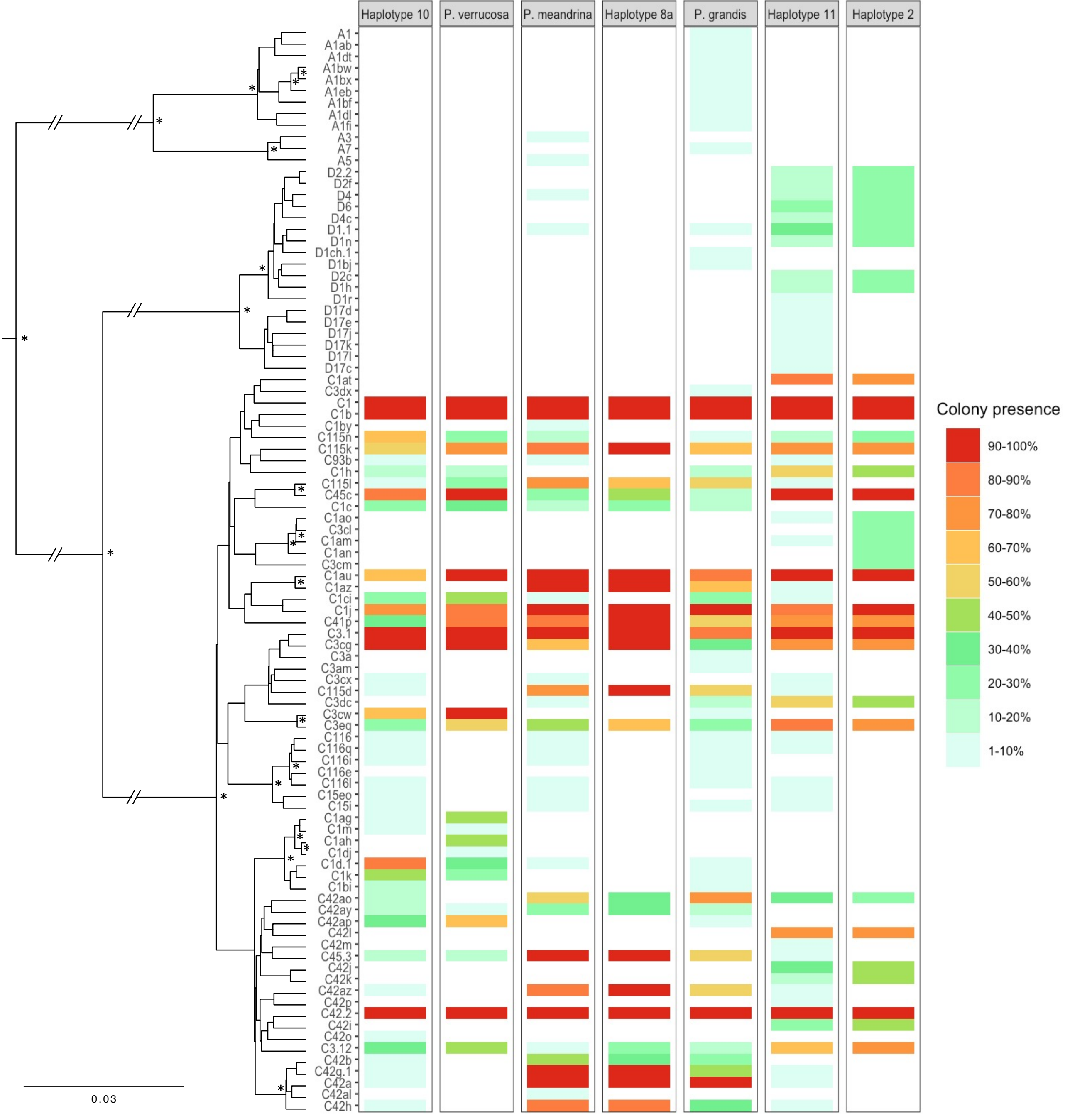

### Supplemental Figure 2

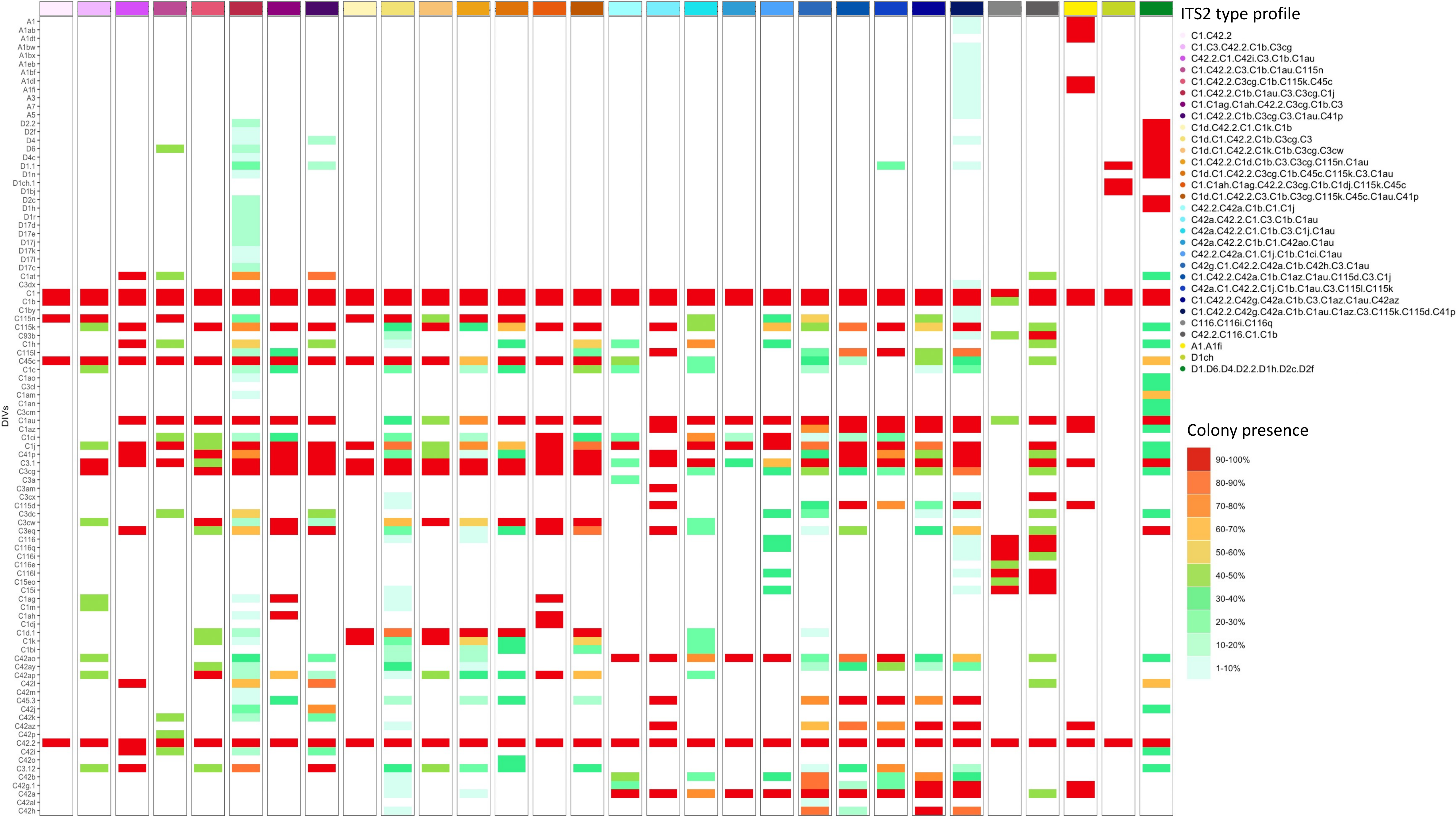
