## Supplemental Figure 3 for "Cophylogeny and specificity between cryptic coral species (*Pocillopora* spp.) at Mo’orea and their symbionts (Symbiodiniaceae)"

### ITS2 Sequences

A1fi
A1
A1ab
A1dl
X1392361_A
X1392372_A
X32746_A
X55778_A
X1042_A
X1362682_A
X759005_A
A1dt
X1392373_A
X1392374_A
X1392363_A
X149403_A
X5706_A
X1392362_A
X1392392_A
X1392391_A
X1392375_A
A7
X140708_A
X1392390_A
X74034_A
X30814_A
X1392364_A
X1392394_A
A5
A1bf
X1392393_A
A1bw
A4
X1392395_A
X1392399_A
A4.3
X1392400_A
X6984_A
X1392396_A
X1892_A
X7747_A
X7695_A
X1392377_A
X1392376_A
A1bx
X7581_A
A3
A1eb
A1fh
A1er
A4a
X73683_A
X759668_A
B1
B5aj
X1390196_B
X29481_B
C1
C42.2
C1b
C3.1
C1d.1
C1au
C42a
C42g.1
C1j
C3cg
C1az
C45c
C115k
C115n
C42h
C41p
X196_C
C42az
C45.3
C42ao
X7_C
C3cw
C116
C115d
C3eq
X93_C
X21709_C
X213_C
X139_C
X566_C
C115l
X114_C
C42b
X526576_C
X348_C
C15
C42l
X99_C
C42ap
X344_C
C3.12
X227_C
C1at
C42ay
C1ag
C1h
C1c
C1k
C1ci
X187_C
C42j
X1936_C
C1m
X1392356_C
X307_C
X312_C
C1ah
X98_C
X33741_C
X278_C
X201_C
C3dc
X317_C
X356_C
X285_C
X287_C
X224_C
C116i
X761_C
X657_C
C15i
X33112_C
X381_C
X197_C
X656_C
X268_C
C116l
X695_C
X192_C
X11_C
X26971_C
X92729_C
C42k
X8128_C
X33398_C
X349_C
X33135_C
X359_C
X985_C
C116q
X8_C
X2790_C
X33434_C
X91390_C
X324_C
X1392365_C
X360_C
X110_C
X234_C
X91056_C
X351_C
X1112172_C
X47161_C
X200_C
C93b
X230_C
X279_C
X138_C
X128_C
X376_C
X21844_C
X101_C
C1bi
X21885_C
X27542_C
C15eo
X119_C
X751_C
X13928_C
X2694_C
X205_C
C1bp
X391641_C
X366_C
X102_C
X202_C
X132_C
X1208585_C
X203_C
X375_C
X21719_C
X355_C
X323_C
X235_C
X653_C
C44
X16816_C
X22186_C
X17_C
X512_C
X1392370_C
X198_C
X5429_C
X62530_C
X33346_C
C42i
C3cx
X316_C
X87_C
X1392402_C
X33522_C
X269_C
X70133_C
X639_C
X6124_C
X284_C
X315_C
X33657_C
X193_C
X273_C
X572_C
X1392403_C
X92006_C
X6125_C
X94456_C
X33327_C
X91839_C
X7844_C
X32893_C
X33249_C
X7305_C
X33703_C
X3246_C
X2193_C
X377_C
X33738_C
X90988_C
X204_C
X662_C
X33388_C
X90755_C
X62136_C
X228_C
X1392369_C
C1dj
X280_C
X33322_C
X47215_C
X6145_C
X1392398_C
X33134_C
X567_C
X33339_C
X327_C
X95383_C
X792_C
X121_C
X43307_C
X33646_C
X206_C
X13271_C
X126_C
X374_C
X372_C
X1392397_C
X68465_C
X346_C
X309_C
X2092_C
X649_C
X68551_C
X33338_C
X33809_C
X90904_C
X10117_C
X380_C
X22_C
X911543_C
X286_C
X694_C
X191_C
C15ev
X33062_C
X222_C
X1274976_C
X281_C
X516_C
X32968_C
X33400_C
X68427_C
X28316_C
X2504_C
C42p
X582058_C
X511_C
X33319_C
X357_C
X89_C
X21705_C
X199_C
C42al
X23144_C
X92079_C
X22172_C
X271_C
X3390_C
X33743_C
X17505_C
X28311_C
X911420_C
X19815_C
X92047_C
X43278_C
X12844_C
X350_C
X2842_C
X196116_C
X33201_C
X386_C
X502415_C
X341_C
X28135_C
X864_C
X483605_C
C15dt
X1211613_C
X276_C
X282_C
X644346_C
X233_C
X90818_C
X1392359_C
X24219_C
X457635_C
C1ca
X1392367_C
X52534_C
X90970_C
X91080_C
X92_C
X270629_C
X23143_C
X15358_C
X651_C
X21721_C
X124_C
X2255_C
X33770_C
X33386_C
X33736_C
X91063_C
X384_C
X33291_C
C93a
X339599_C
X66892_C
X14578_C
X476678_C
X90864_C
X22181_C
X33435_C
X486744_C
X91113_C
X100_C
X18628_C
X33343_C
X91244_C
X137_C
X911410_C
X23486_C
X33565_C
X52533_C
X595337_C
X8872_C
X1392404_C
X913214_C
X11391_C
X21507_C
X26594_C
X5403_C
X52535_C
X90577_C
X658403_C
X1215183_C
X33740_C
X33122_C
X479399_C
X460980_C
X42931_C
X21813_C
X91428_C
X33802_C
X21_C
X911416_C
X208_C
X14278_C
X1392357_C
X1392406_C
X10943_C
X33663_C
X95517_C
X433981_C
X1392378_C
C15bc
X271833_C
X749572_C
X911417_C
X502403_C
X91288_C
X10568_C
X36147_C
X33120_C
X658970_C
X8117_C
X58726_C
X1392401_C
X68496_C
X33268_C
X90753_C
C3a
X392_C
X911409_C
X209_C
X476707_C
X91064_C
X532307_C
X27636_C
X33072_C
X771_C
C1am
X6157_C
X435_C
X86_C
X26674_C
X11683_C
X36151_C
X33309_C
X91074_C
X212679_C
X14791_C
X911419_C
X483586_C
X49033_C
C42m
X278299_C
X1318267_C
X770_C
X220_C
X91180_C
X8371_C
C1w
C1r
X95042_C
X751238_C
X270_C
X43704_C
X95278_C
X962718_C
X90_C
X911421_C
X33733_C
X21714_C
X911424_C
X502494_C
X28244_C
X557_C
X26682_C
X91806_C
C42ax
C3.2
X553_C
X92781_C
X1211131_C
X13421_C
C42o
X95088_C
X47223_C
X91240_C
X90862_C
C1by
X336707_C
X10283_C
X404696_C
X33070_C
X33121_C
X90622_C
X40148_C
X21931_C
X2864_C
X52541_C
X62554_C
X12576_C
X1271393_C
X466820_C
X13806_C
X26871_C
X33433_C
X93211_C
X13671_C
C3am
X90534_C
C1ao
X90897_C
X33401_C
X1392371_C
X33608_C
X758_C
X1392405_C
X90861_C
X96427_C
X93045_C
X194_C
X5355_C
X816_C
X497295_C
X519504_C
X1355_C
X1392366_C
C116e
X911437_C
X396_C
X207190_C
X33803_C
X90560_C
X91274_C
X26679_C
C3dx
X456567_C
X911422_C
X500632_C
X21747_C
X32888_C
X911515_C
X91296_C
X26204_C
X92652_C
X71616_C
X90618_C
X90652_C
X33348_C
X90849_C
X21547_C
X765_C
X1392358_C
X97_C
X911476_C
X114220_C
X911438_C
X90629_C
X911540_C
X545310_C
X1392368_C
X842_C
X91087_C
X21804_C
X33399_C
X465500_C
X911468_C
X911547_C
X519527_C
X22296_C
X61826_C
C1an
X1392360_C
X61822_C
C3cm
X90811_C
X33385_C
X23115_C
X27927_C
C3cl
X7532_C
C1x
D1.1
D6
D4
D1ch.1
D2.2
D2f
X763946_D
X1760_D
D1h
D2c
X14166_D
D1bj
X439757_D
X132119_D
X10082_D
X16303_D
X1392379_D
X447436_D
X446428_D
D4c
X440001_D
D1n
X61933_D
X25432_D
X1392380_D
X1392381_D
D17d
X1392382_D
X1392384_D
X1392383_D
X15842_D
D17e
X1392385_D
X1392386_D
X1392387_D
X10614_D
D17c
X9233_D
X1392388_D
D17j
D1r
D17l
X12538_D
X10616_D
D17k
X260537_D
X10612_D
X1392389_D
X946_D
X7280_D
X19420_D
X179669_D
